# Activity labeling *in vivo* using CaMPARI2 reveals electrophysiological differences between neurons with high and low firing rate set points

**DOI:** 10.1101/795252

**Authors:** Nicholas F Trojanowski, Gina G. Turrigiano

## Abstract

Individual excitatory neurons in visual cortex (V1) display remarkably stable mean firing rates over many days, even though these rates can differ by several orders of magnitude between neurons. When perturbed, each neuron’s firing rate is slowly regulated back to its pre-perturbation level, demonstrating that neurons maintain their mean firing rate around an individual firing rate set point (FRSP). To better understand the mechanisms that neurons within a single cell type use to maintain different FRSPs *in vivo*, we implemented a novel method of activity labeling that uses CaMPARI2, a fluorescent protein that undergoes Ca^2+^- and UV-dependent green-to-red photoconversion, to permanently label neurons in freely behaving mice based on their firing rates. We found that immediate early gene (IEG) expression was correlated with CaMPARI2 red/green ratio following an activity stimulation paradigm, and that neurons with greater photoconversion *in vivo* tended to have a higher firing rate *ex vivo*. In layer 4 (L4) pyramidal neurons in mouse monocular V1, which comprise a single transcriptional cell type, we found that high activity neurons had a left-shifted F-I curve, lower rheobase current, and decreased spike adaptation index relative to low activity neurons, demonstrating increased intrinsic excitability. Surprisingly, we found no difference in total excitatory or inhibitory synaptic current or in E/I ratio between high and low activity neurons. Thus, within a single cell type differences in intrinsic excitability and spike frequency adaptation can contribute to divergent activity set points. These results reveal that E/I ratio plays only a minor role in determining the firing rate set point of L4 pyramidal neurons, while intrinsic excitability is an important factor.

## Introduction

Decades of research on different forms of homeostatic plasticity have convincingly demonstrated that neuronal circuit activity is tightly regulated despite dynamic inputs (Davis, 2013; Marder, 2011; Turrigiano, 2008). This regulation is essential for maximizing information storage and transfer, and for preventing hypo- or hyper-excitability (Abbott and Nelson, 2000; Turrigiano, 2011). In vertebrates, homeostatic plasticity is perhaps best understood in the somatosensory and visual cortex, where it has become clear that neuronal activity can be homeostatically stabilized by either isolated or concurrent changes in synaptic strength and intrinsic excitability (Gainey and Feldman, 2017). Disruptions in homeostatic plasticity have been proposed to contribute to a wide range of neurological disorders, including Alzheimer’s disease (Styr and Slutsky, 2018), epilepsy, and autism spectrum disorders (Ebert and Greenberg, 2013; Nelson and Valakh, 2015), suggesting that understanding how homeostatic set points are built and maintained will generate important insight into a range of circuit dysfunctions.

Pyramidal neurons in rodent cortex exhibit remarkably stable firing rates over hours to days, even though these firing rates can span multiple orders of magnitude across neurons (Dhawale et al., 2017; Hengen et al., 2016; Keck et al., 2013; Torrado Pacheco et al., 2019). In response to prolonged sensory deprivation, pyramidal neurons initially decrease their firing but eventually homeostatically return to their own individual firing rate set point (FRSP), a process driven by changes in synaptic strength and intrinsic excitability (Hengen et al., 2013; Lambo and Turrigiano, 2013; Maffei et al., 2004; Nataraj and Turrigiano, 2011). High and low FRSP neurons in rodent visual cortex and hippocampus are differentially modulated by sleep states, suggesting that high and low FRSP neurons are functionally distinct (Miyawaki et al., 2019; Miyawaki and Diba, 2016; Watson et al., 2016), but little is known about the specific electrophysiological differences that generate these differences in FRSP.

The mechanisms that control FRSPs *in vivo* have not been studied in any brain region, in part due to the difficulty of recording the activity of individual neurons *in vivo*, then re-identifying the same neurons *ex vivo* for further examination. Targeted approaches for labeling individual neurons based on their *in vivo* responses are low throughput and prone to sampling bias (Gilbert and Wiesel, 1979; Lien and Scanziani, 2011; Pinault, 1996). Reconstruction of serial sections following *in vivo* Ca^2+^ imaging is possible but requires unambiguous realignment of acute slices with *in vivo* images (Ko et al., 2011). Labeling neurons based on immediate early gene (IEG) expression can identify active subsets of cells (Yassin et al., 2010), but the relationship between IEG expression and firing rate is not linear (Tyssowski and Gray, 2019).

To label neurons *in vivo* based on their FRSP we used CaMPARI2, a fluorescent mEos derivative that undergoes Ca^2+^- and UV-dependent photoconversion from green to red (Fosque et al., 2015; Moeyaert et al., 2018; Zolnik et al., 2016). Using low illumination power and long duration photoconversion (15 to 30 min), we were able to label neurons in monocular visual cortex (V1m) based on their mean firing rates in freely behaving mice. Following a paradigm that transiently increases neuronal activity, we found a correlation between cFos expression and *in vivo* CaMPARI2 photoconversion. We found that the red/green ratio of individual neurons was lognormally distributed, as were firing rates when recorded *ex vivo*, and neurons with greater *in vivo* photoconversion tended to have higher firing rates *ex vivo*. In layer 4 (L4) pyramidal neurons, we found that high activity neurons had a greater intrinsic excitability than low activity neurons, though we found no evidence of differences in excitatory or inhibitory charge or in excitation/inhibition (E/I) ratio. These data demonstrate that CaMPARI2 can be used to label cells based on their activity *in vivo*, and suggest that intrinsic excitability is an important determinant of FRSP.

## Results

### CaMPARI2 photoconversion *in vivo* labels neurons based on their activity levels

In order to test if CaMPARI2 could be used as a permanent marker of *in vivo* activity, we injected an AAV containing CaMPARI2 into V1m of mice at postnatal day 15 (P15). One week later, we implanted a fiberoptic in the same craniotomy hole used for virus injection. One to two weeks after cannula implantation, mice were subjected to *in vivo* photoconversion. Brief (1 to 2 s) photoconversion of CaMPARI2 has previously been used to identify neurons that respond to a specific sensory stimulus (Moeyaert et al., 2018), but here we sought to label neurons based on their mean firing rates in freely behaving and viewing animals, so longer stimulation times were required. Photoconversion was performed during the light cycle, when animals were freely behaving. We found that with low UV light power (~0.20 mW), 15 to 30 minutes of UV illumination was sufficient to cause robust photoconversion without excessive tissue damage. Due to the light-scattering properties of brain tissue, light intensity attenuates with increasing distance from the source, especially at shorter wavelengths (Al-Juboori et al., 2013). We found that the average red/green CaMPARI2 photoconversion ratio was quite constant within 240 μm of the fiberoptic cannula (Fig. 1A), so all subsequent experiments were performed on neurons within this distance.

**Figure 1.**
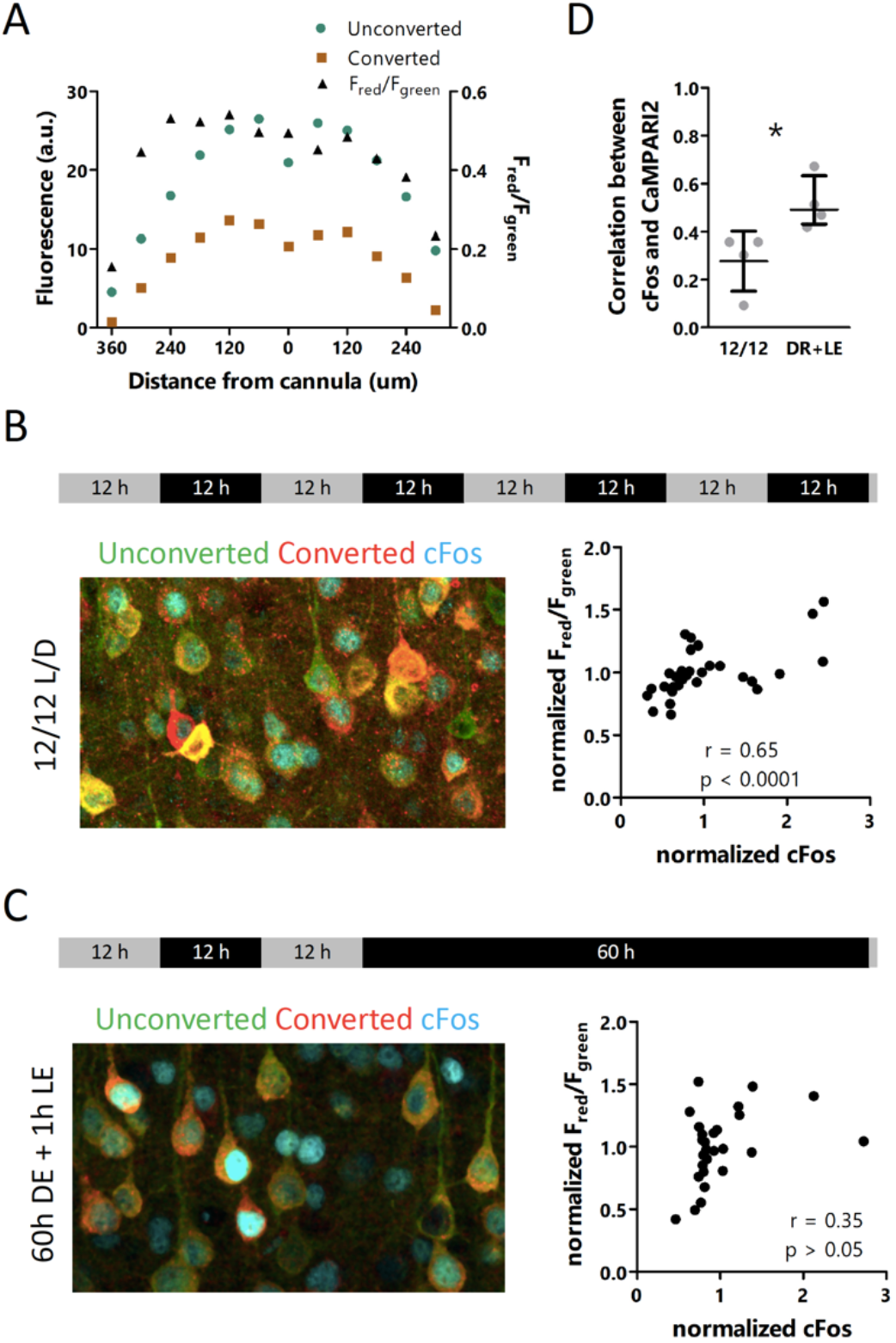
CaMPARI2 photoconversion *in vivo* is correlated with activity-dependent cFos levels. **A:** CaMPARI2 red and green fluorescence values (left axis) and red/green ratio (right axis) as a function of distance from the fiberoptic cannula. **B:** Experimental paradigm (top), immunofluorescence of cFos (cyan) and red and green forms of CaMPARI2 for mice housed in conventional 12/12 light/dark housing (left), and correlation between red/green ratios and cFos expression (right). n = 31 cells, r = Spearman’s rank correlation coefficient. **C:** Experimental paradigm (top), immunofluorescence of cFos (cyan) and red and green forms of CaMPARI2 for mice subjected to 60 hours of dark exposure followed by 1 hour of light re-exposure (left), and correlation between red/green ratios and cFos expression (right). n = 27 cells. r = Spearman’s rank correlation coefficient. **D:** Correlations between red/green ratios and cFos expression for control and elevated firing conditions. Bars represent mean +/− 95% CI; n = 4 animals/condition, 23 to 45 cells/animal; * p<0.05, Mann Whitney U test.

To test if neurons with a higher red/green CaMPARI2 ratio after *in vivo* photoconversion represented neurons with increased activity, we first compared this ratio to an established marker of neuronal activity. Labeling with IEGs such as cFos is a common approach for identifying neurons that have recently undergone activity-dependent gene transcription (Barth et al., 2004; Sagar et al., 1988; Yap and Greenberg, 2018). Recent work in the rodent visual cortex has revealed that 60 hours of dark exposure followed by 1-hour light re-exposure, a protocol known to cause robust expression of IEGs including cFos, also causes elevated firing rates for 15 to 30 minutes after lights on (Torrado Pacheco et al., 2019). Therefore, we exposed mice to this protocol while photoconverting CaMPARI2 during the first 30 minutes of light re-exposure. In contrast to mice housed on a normal 12/12 light/dark cycle, where we saw little correlation between the two signals (Fig. 1B), after light re-exposure we saw a significant correlation between cFos levels and *in vivo* CaMPARI2 photoconversion red/green ratio (Fig. 1C, D). This is consistent with previous work that showed that cFos-positive neurons had higher activity than cFos negative neurons (Yassin et al., 2010), and suggests that *in vivo* photoconversion of CaMPARI2 can differentiate between high and low activity neurons.

To determine if neurons with different firing rates *in vivo* retained these properties in acute slices, we used cell-attached recordings from acute slices in active ACSF to measure the activity of excitatory neurons previously photoconverted *in vivo* (Fig. 2A-C). We found that the distribution of firing rates was lognormal and very similar to the distribution observed from chronic recordings *in vivo* (Fig. 2D) (Hengen et al., 2016). The distribution of red/green ratios was also lognormal, though over a slightly narrower range than the firing rate distribution (Fig. 2E). Taken together, these observations indicate that many of the features that generate a lognormal firing rate distribution *in vivo* are conserved in the active slice preparation, and that CaMPaRI2 photoconversion is able to capture a similar range of variation in activity.

**Figure 2:**
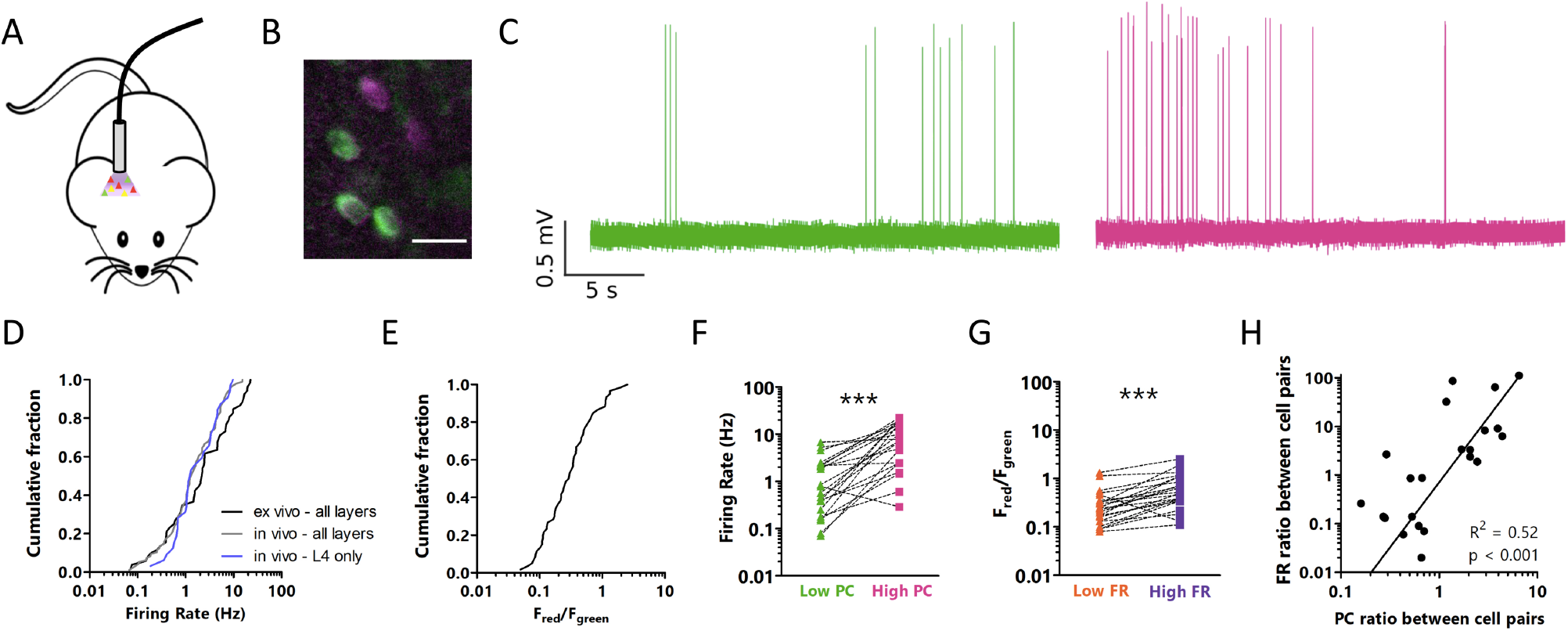
CaMPARI2 photoconversion *in vivo* labels neurons based on their activity levels. **A**: Cartoon representation of *in vivo* CaMPARI2 photoconversion. **B**: Representative *ex vivo* image of *in vivo* CaMPARI2 photoconversion; **C**: Representative cell-attached recordings from neurons with low (left) and high (right) *in vivo* photoconversion. **D**: Cumulative fraction of firing rates of putative excitatory neurons measured in acute slices in active ACSF, or from chronic *in vivo* recordings from all layers or only from electrodes positioned in L4. n = 52 cells *ex vivo*, 96 cells *in vivo* all layers, 32 cells *in vivo* L4. **E**: Cumulative fraction of red/green ratios for putative excitatory neurons. Neurons were photoconverted *in vivo*, and red/green ratio was measured in acute slices. n = 52 cells. **F**: Difference between *ex vivo* firing rates for pairs of excitatory neurons photoconverted *in vivo*. For each pair, the neuron with the higher red/green ratio was designated the High PC neuron. **G**: Difference between red/green ratios of higher and lower FR neurons. Neurons were photoconverted in vivo, and firing rates were measured *ex vivo* in active slice. **H**: Correlation between the difference in *in vivo* red/green ratios and *ex vivo* FR ratios between pairs of neurons. Line represents linear best fit. F, G, H: n = 22 pairs from 11 animals; *** p<0.001, paired t-test. D through H, neurons were sampled from all layers.

Next, we wished to verify that high *in vivo* photoconversion neurons have higher firing rates *ex vivo* than low photoconversion neurons. Since CaMPARI2 photoconversion rate depends on UV intensity (Moeyaert et al., 2018), the red/green CaMPARI2 photoconversion ratio can best be compared between neurons that have received a similar amount of light. Therefore, for this and all subsequent experiments (except where noted), we recorded from pairs of nearby neurons with different red/green photoconversion ratios. Within neuron pairs, in almost all cases we found that the neuron that had a higher red/green ratio (indicating greater *in vivo* activity) also had a greater firing rate *ex vivo* (Fig. 2F). When we sorted the neurons within each pair according to *ex vivo* firing rate, in most cases the higher firing rate cell also had a higher red/green ratio (Fig. 2G). The magnitude of differences in firing rate and red/green ratio in each slice were correlated, such that neurons that had a large difference in firing rate tended to also have a large difference in red/green ratio, and vice versa (Fig. 2H). Thus, many of the factors that lead to a difference in firing rate between neurons *in vivo* are preserved in the acute slice preparation.

### High activity pyramidal neurons in L4 have greater intrinsic excitability than low activity neurons

Differences in intrinsic excitability and synaptic connectivity have been found between different cell types in L5 of visual cortex (Hattox and Nelson, 2007; Kim et al., 2015), but it is unknown what factors generate high or low FRSPs within a single cell type. To begin to answer this question, we measured the intrinsic excitability and synaptic inputs of excitatory neurons in L4, since these neurons cluster into a single transcriptional cell type (Tasic et al., 2018), yet have a broad distribution of mean firing rates (Fig. 2d).

Using the paired approach described above, in acute slices we measured the intrinsic excitability of *in vivo* photoconverted high and low activity star pyramidal neurons in L4 by giving a series of current step injections in the presence of synaptic blockers (Fig. 3A-C). By comparing the areas under the firing rate-current (F-I) plot, we found that high activity neurons in L4 have a greater intrinsic excitability than low activity neurons (Fig. 3D, E). This difference can be partially attributed to decreased rheobase current in high activity neurons (Fig. 3F), though even when firing rates were measured relative to rheobase a difference in intrinsic excitability was still present (Fig. 3G, H). This rheobase-independent difference in excitability was accompanied by a difference in spike frequency adaptation: low activity neurons showed a greater degree of adaptation, and thus fewer spikes for a given current step, than high activity neurons (Fig. 3I). This effect was present regardless of how spike trains were selected for comparison (Fig. 3J). In contrast to the correlation between input resistance and rheobase, and between input resistance and area under the non-adjusted F-I curve (Fig. S1), we saw no correlation between input resistance and spike frequency adaptation (Fig. 3K), suggesting that there are at least two distinct facets of intrinsic excitability that differ between high and low firing rate neurons. We found no significant differences in other aspects of action potential waveform or kinetics, or in afterhyperpolarization or voltage sag (Table S1).

**Figure 3:**
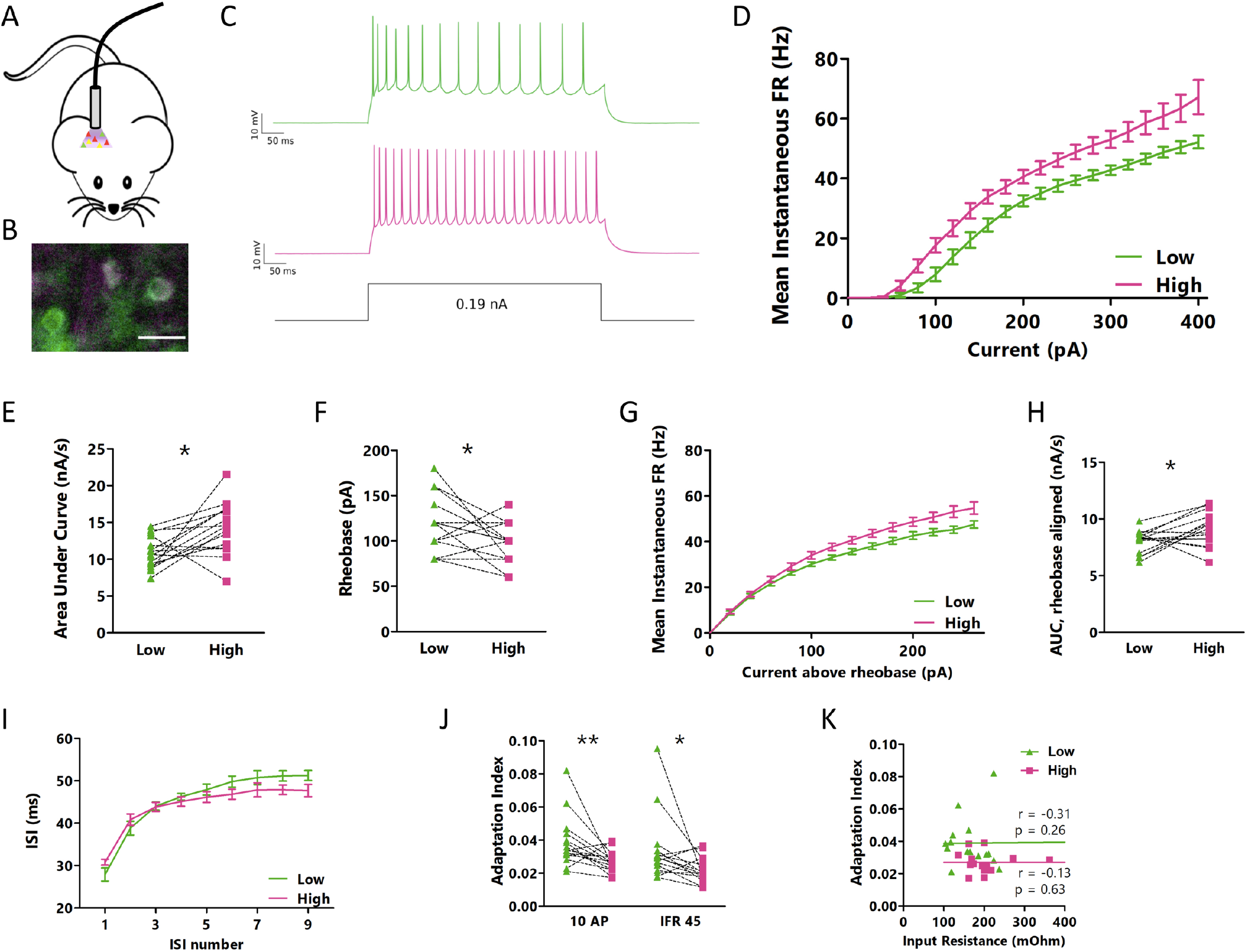
High and low activity pyramidal neurons in L4 differ in multiple aspects of intrinsic excitability. **A**: Cartoon representation of *in vivo* CaMPARI2 photoconversion. **B**: Representative *ex vivo* image of *in vivo* CaMPARI2 photoconversion. **C**: Representative current clamp recordings for intrinsic excitability measurements. **D**: Mean instantaneous firing rate versus current injection (F-I) for pairs of neurons photoconverted *in vivo*. **E**: Comparison of the area under the F-I curve (from D) for each neuron pair. **F**: Comparison of the rheobase values for each neuron pair. **G**: Rheobase-subtracted mean instantaneous F-I curve for neuron pairs. **H**: Comparison of the area under the curves in G for each neuron pair. **I**: Inter-spike interval versus action potential number for the smallest current step that produced at least 10 action potentials for each neuron pair. **J**: Adaptation index for high and low FR neurons. Traces were selected from different current steps depending on the number of action potentials (left, smallest current injection to produce 10 action potentials) or mean instantaneous firing rate (right, smallest current injection to produce a mean instantaneous firing rate of 45 Hz). **K**: Correlation between input resistance and adaptation index (for 10 AP current step) for all neurons. D through K: n = 15 pairs of cells from 7 animals; * p<0.05, ** p<0.05, E, F, H: paired t-test; H, Wilcoxon matched-pairs signed rank test; J: r = Spearman’s rank correlation coefficient. All recordings and images are from pyramidal neurons in L4 V1.

If differences in intrinsic excitability are important for the differences in FRSP *in vivo*, and activity profiles *ex vivo* accurately reflect *in vivo* activity, then neurons with higher *ex vivo* firing rates should also have greater intrinsic excitability. To test this, we devised an *ex vivo* photoconversion protocol in active acute slices (Fig. 4A, B, see Methods for details). This approach has the advantage that illumination is uniform in the X-Y plane so we are not limited to the paired approach, and thus can directly compare several neurons from a given slice. After 30 minutes of *ex vivo* photoconversion in the active slice preparation, we measured the intrinsic excitability of 4 to 6 neurons per slice. We found a strong correlation between greater photoconversion and higher intrinsic excitability (Fig. 4C, D). Thus, differences in intrinsic excitability appear to play an important role in determining each neuron’s mean activity level.

**Figure 4:**
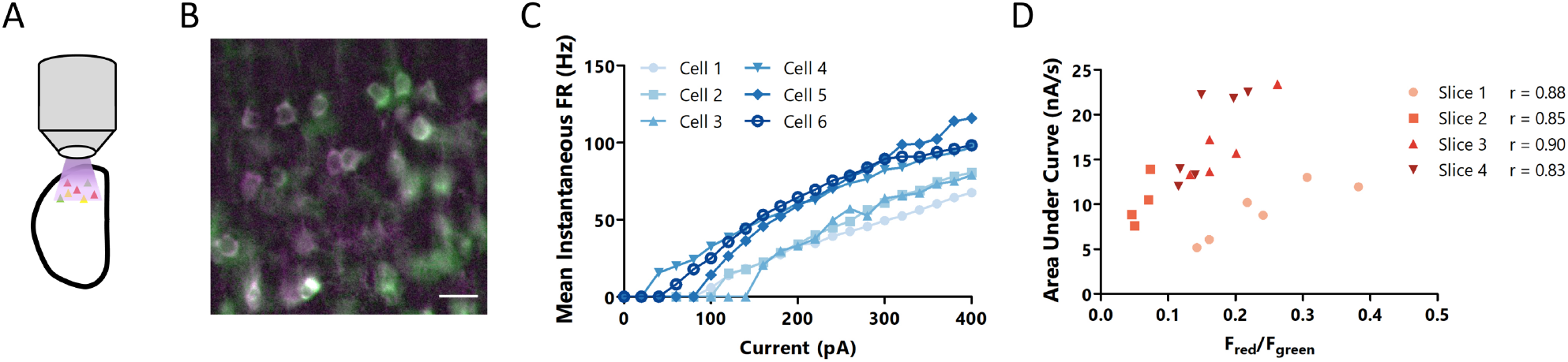
L4 pyramidal neurons with higher activity *ex vivo* have greater intrinsic excitability than neurons with lower activity. **A**: Cartoon representation of *ex vivo* CaMPARI2 photoconversion. **B**: Representative *ex vivo* image of *ex vivo* CaMPARI2 photoconversion. **C**: Mean instantaneous firing rate versus current injection for six neurons from the same slice, photoconverted *ex vivo*. Darker colors represent a greater red/green ratio. **D**: Correlation between the area under the F-I curve and the red/green ratio following *ex vivo* photoconversion. Each point represents one neuron, and each slice is plotted in the same color. n = 4 to 6 cells per slice. r = Spearman’s rank correlation coefficient. All recordings and images are from pyramidal neurons in L4 V1.

### High and low activity excitatory neurons in L4 of visual cortex receive similar net synaptic excitation and inhibition

Having established a robust difference in intrinsic excitability between high and low activity neurons, we next sought to determine if these neurons differed in the amount of synaptic input they received. To measure the relative total excitatory and inhibitory drive onto high and low firing rate neurons, we used voltage clamp to hold neurons at the experimentally determined reversal potential for either inhibition or excitation (typically −60 mV and +5 mV, respectively) in an active slice preparation, while blocking action potentials intracellularly. This allowed us to sequentially measure the total excitatory and inhibitory synaptic charge onto individual neurons (Fig. 5A-C). Strikingly, there were no consistent differences in either excitatory or inhibitory charge between high and low activity neurons. We also found that although the ratio of excitatory to inhibitory charge was broadly distributed across neurons, there was no correlation between this ratio and whether the neuron had a high or low mean firing rate (Fig. 5D, E). Likewise, there was no difference in the frequency of excitatory or inhibitory potentials between high and low activity neurons (Fig. 5F, G). While these results do not exhaustively rule out a role for synaptic input in determining the FRSP, they do support a model where intrinsic excitability is a more important determinant of FRSP than E/I ratio.

**Figure 5:**
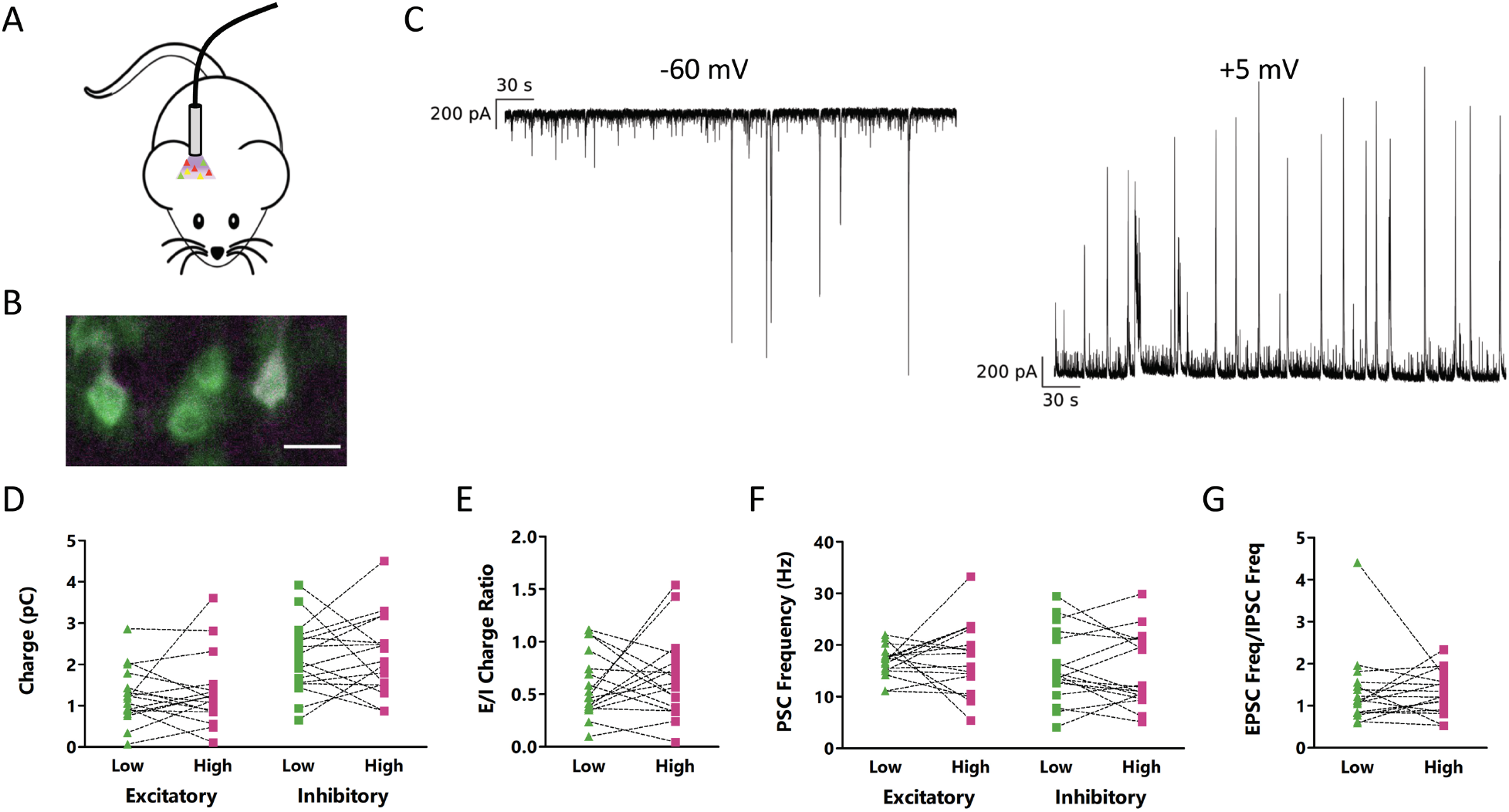
High and low activity L4 pyramidal neurons have no consistent differences in excitatory or inhibitory synaptic drive, or E/I ratio. **A**: Cartoon representation of *in vivo* CaMPARI2 photoconversion. **B**: Representative *ex vivo* image of *in vivo* CaMPARI2 photoconversion. **C**: Representative traces showing all excitatory (left) and inhibitory (right) current. To isolate the input type, recordings were taken at the reversal potential for inhibition (left) or excitation (right) in active ACSF in the absence of synaptic blockers. **D**: Total excitatory (left) or inhibitory (right) synaptic charge measured in pairs of neurons at their experimentally determined inhibitory (left) or excitatory (right) reversal potential. **E**: Ratio of total excitatory to inhibitory charge for pairs of photoconverted neurons. **F**: Frequency of excitatory (left) or inhibitory (right) synaptic currents measured in pairs of neurons at their experimentally determined inhibitory (left) or excitatory (right) reversal potential. **G**: Ratio of frequency of excitatory to inhibitory currents for pairs of photoconverted neurons. All recordings and images are from pyramidal neurons in L4 V1. D through G: n = 17 pairs of cells from 11 animals. No significant differences, paired t-test.

## Discussion

Mean firing rates of neocortical neurons *in vivo* span many orders of magnitude and can be stable over days (Dhawale et al., 2017; Hengen et al., 2016), but the factors that determine where a given neuron sits in this broad firing rate distribution are not understood. Using the photoconvertible activity marker CaMPARI2, we were able to permanently label neurons based on their mean activity levels *in viv*o in freely behaving mice. This powerful approach allowed us to explore the differences that underlie the broad, lognormal distribution of firing rates that is a ubiquitous feature of neuronal activity in many brain regions (Buzsáki and Mizuseki, 2014). Investigation of the electrophysiological differences between high and low activity pyramidal neurons in L4 of mouse visual cortex – considered a single molecular cell type – revealed that high activity neurons have greater intrinsic excitability than low activity neurons, whereas excitation/inhibition ratio is not consistently different between high and low activity neurons. Together, these results argue that the intrinsic excitability of a neuron, which is determined by the interaction of a diverse array of membrane currents, controls a feature of cellular identity (firing rate set point) that is orthogonal to the standard transcriptional definition of a cell type.

For identifying neurons with different FRSPs, CaMPARI2 has a number of advantages over other common methods of activity-labeling. First, the mark is permanent and is preserved in *ex vivo* preparations, unlike conventional calcium indicators. Second, although immediate early genes (IEGs) such as cFos have been used for decades to identify neurons that have been recently active (Yap and Greenberg, 2018), IEG expression is not a linear readout of activity and the exact relationship between IEG expression levels and neuronal activity is unclear (Sheng et al., 1993; Tyssowski and Gray, 2019). Further, cFos expression seems to better capture changes in – rather than absolute levels of – activity, as suggested by our observation that there is little correlation between cFos and CAMPARI2 signals during normal activity, but a correlation emerges when firing is transiently elevated through dark-exposure followed by re-exposure to light. In contrast to the thresholding effect of the cFos signal, we find a broad and lognormal distribution of CAMPARI2 *in vivo* photoconversion rates that closely mirrors the distribution of *in vivo* firing rates, suggesting that CAMPARI2 photoconversion ratios capture much of the variation in firing rates seen *in vivo*.

There are a few factors that could confound the use of CaMPARI2 for activity labeling, most of which are shared with rapid Ca^2+^ indicators such as GCaMP6s. First, its photoconversion ratio is dependent on Ca^2+^, rather than action potentials, so any difference in Ca^2+^ buffering or in synapse-mediated calcium influx between neurons will make it difficult to accurately measure their relative firing rates. Despite this potential caveat, within pairs of neurons we were able to show a strong correlation between *in vivo* photoconversion of CaMPARI2 and *ex vivo* spontaneous firing rates. Second, since the photoconversion rate of CaMPARI2 depends on the amount of UV light received, it is challenging to use CaMPARI2 to examine differences in activity over a wide area *in vivo*. Improvements in cannula design (Pisanello et al., 2017) may mitigate these issues somewhat, though this will likely remain a fundamental problem with CaMPARI2 use *in vivo*, where UV transmittance is poor. The use of paired recordings from nearby neurons with distinct red/green ratios allowed us to circumvent this limitation, and the agreement between our *in vivo* and *ex vivo* data provides confidence that the differences we observe following either photoconversion protocol capture meaningful differences in neuronal activity.

It has been broadly postulated that a major contributor to neuronal excitability is the ratio of excitation to inhibition a neuron receives (Isaacson and Scanziani, 2011), so we were surprised to find no systematic difference in E/I charge ratio for low and high firing rate neurons. One possible interpretation of this finding is that important differences in synaptic drive are missing in our *ex vivo* preparation, which preserves local connectivity but not longer-range connections. While we cannot entirely rule this out, a few considerations suggest that this is unlikely to be a major factor. First and most importantly, we find a very similar lognormal firing rate distribution in our active *ex vivo* preparation as we find *in vivo*. Second, within pairs of neurons we find a strong correlation between *in vivo* photoconversion and *ex vivo* firing rates. Taken together, these two observations suggest that many of the factors that generate this broad firing rate distribution are preserved between the two preparations. The most striking difference we find between low and high firing rate L4 pyramidal neurons is a constellation of differences in intrinsic excitability, which likely involve differential expression of a variety of voltage-dependent ion channels. This difference in intrinsic excitability is well-correlated with the difference in CaMPARI2 photoconversion, but it is not clear that the range of intrinsic excitability differences we observe entirely accounts for the wide range of mean firing rates. There may well be other features of synaptic inputs aside from total charge that also contribute to this broad distribution, such as the identity or the temporal characteristic of neuronal inputs. Nonetheless, our data show that a major determinant of differences in firing between L4 pyramidal neurons is differences in intrinsic excitability.

In addition to firing rates, higher order features of V1 activity such as the coefficient of variation of inter-spike interval, pairwise spike correlation structure, and criticality are also stable over time and under homeostatic control (Hengen et al., 2016; Ma et al., 2018; Wu et al., 2019). Modeling work suggests that distinct homeostatic plasticity mechanisms control these different features of network function. For example, synaptic scaling can serve to preserve the correlation structure of local networks, while intrinsic homeostatic plasticity is well-suited to restore firing rates after perturbations (Wu et al., 2019). While some conserved features of network connectivity may require non-cell-autonomous regulatory mechanisms (Ma et al., 2018), our data suggest that FRSP is largely determined through cell-autonomous differences in intrinsic excitability.

There are theoretical limits on neuronal firing rates due to energetic considerations. It has been estimated that average firing rates across the human brain cannot exceed 0.94 Hz without generating an unmeetable energy burden (Lennie, 2003), though this value is likely somewhat higher for the rodent brain (Attwell and Laughlin, 2001; Howarth et al., 2012). Data from cultured hippocampal neurons suggest that mitochondrial metabolism plays an important role in constraining the ensemble firing rate set point (Frere and Slutsky, 2018; Styr et al., 2019). Consistent with our data, reducing mitochondrial spare respiratory capacity reduces both the ensemble FRSP and the intrinsic excitability of hippocampal neurons (Styr et al., 2019). The molecular mechanisms that link mitochondrial function to intrinsic excitability are unclear, as is how these metabolic limits on firing allow individual neurons to express such a broad range of FRSPs. It is possible that stochastic differences in mitochondrial function across neurons causes differences in intrinsic excitability and FRSP to emerge, but this idea has yet to be tested.

Recently, there has been a strong push to catalog and sort all of the neurons in various brain regions, or even the entire brain (Economo et al., 2018; Gouwens et al., 2019; Phillips et al., 2019; Saunders et al., 2018; Sugino et al., 2019; Tasic et al., 2018; Zeisel et al., 2018, 2015), using various approaches to separate cells into classes. Most popular have been techniques that sort cells based on their transcriptional profile or projection targets, though efforts are ongoing to incorporate this anatomical and transcriptional data with electrophysiological data (Gouwens et al., 2019). Given the characteristic lognormal distribution of firing rates across multiple regions of cortex (Buzsáki and Mizuseki, 2014) and the stability of individual neurons within this distribution (Dhawale et al., 2017; Hengen et al., 2016), we argue that FRSP represents a feature of cellular identity that is orthogonal to anatomical or transcriptional cell type. The FRSP of a neuron cannot necessarily be captured by a battery of electrophysiological tests performed *ex vivo*, but rather require labeling neurons based on their mean activity *in vivo*. Activity labeling with CaMPARI2 allows this *in vivo* labeling of neurons followed by *ex vivo* analysis of excitability and connectivity, a crucial step for understanding of the genesis, regulation, and function of FRSPs.

## Materials and Methods

### Mice

Experiments were performed on WT C57BL/6J mice or B6;C3-Tg(Scnn1a-cre)3Aibs/J mice of both sexes (Miska et al., 2018). Experiments were performed between postnatal day (P) 30 and 40 for *in vivo* photoconversion experiments, and between P22 and P25 for *ex vivo* photoconversion experiments. Mice were housed on a 12/12 light/dark cycle unless otherwise indicated. Food and water were available *ad libitum*. All animals were housed, cared for, and sacrificed in accordance with Brandeis IBC and IACAUC protocol.

### Virus construction

pAAV_hSyn1_NES-his-CaMPARI2-F391W-WPRE-SV40 (AAV9), used in Figures 2, 3, and 5, was a gift from Benjamin Moeyaert and Eric Schreiter, Janelia Research Campus.

pAAV_CaMKII_NES-his-CaMPARI2-F391W-WPRE-SV40 (AAV9), used for Figure 1, was made by the replacing the hSyn1 promoter in hSyn1_CaMPARI2 with a 0.4 kb CaMKII promoter fragment (Prakash et al., 2012), then ligating CaMKII-CaMPARI2-WPRE-SV40 into a pAAV-MCS backbone (gift from Yasuyuki Shima and Sacha Nelson, Brandeis University).

pAAV_hSyn1_Flex_NES-his-CaMPARI2-F391W-WPRE-SV40 (AAV9), used for Figure 4, was constructed by a ligating vector inserts containing hSyn1-Flex-CaMPARI2-WPRE-SV40 (gift from B. Moeyaert and E. Schreiter) into the pAAV-hSyn1-Flex-mRuby2-GSG-P2A-GCaMP6s-WPRE-pA backbone in place of the Flex-GCaMP6s-WPRE-pA (Addgene 68720).

### Dark exposure and light re-exposure

Animals were raised normally until P21, then placed into a custom-built light-tight dark box for 60 hours (Torrado Pacheco et al., 2019). After 60 hours, animals were anesthetized with ketamine/xylazine/acepromazine then transcardially perfused either immediately following removal from darkness, or after 1 hour of light re-exposure.

### Surgeries

Stereotaxic viral injections were performed between P15 and P17 under ketamine/xylazine/acepromazine anesthesia (Miska et al., 2018). V1m was targeted using stereotaxic coordinates after adjusting for the measured lambda-bregma distance. A glass pipette pulled to a fine point delivered 200 nL of virus at the target depth through a targeted craniotomy.

Cannula implantation was performed between P22 and P25 under isoflurane anesthesia. Two small machine screws (303 Stainless Steel Machine Screw, Antrin) were inserted into the skull for stability, then fiberoptic cannulas fabricated in-house (core diameter 250μm, cannula diameter 1.25 mm, NA 0.66, length ~1.5 mm, supplies from Prizmatix) were placed through the hole used for virus injection. Dental cement was then used to cover the skull and screws and anchor the cannula in place.

### CaMPARI2 *in vivo* photoconversion

For *in vivo* photoconversion, a fiberoptic cable (Optogenetics Fiber −500-1.25, Prizmatix) was connected to the implanted cannula and mice were transferred to a photoconversion arena containing *ad libitum* food. After 10 minutes of acclimation, the 390 nm light (Silver-LED-390B, Prizmatix) was turned on for 30 minutes at ~0.25 mW.

### Acute slice preparation

For acute slice experiments, mice were deeply anesthetized with isoflurane immediately following photoconversion. After slicing in carbogenated (95% O2, 5% CO2) standard ACSF (in mM: 126 NaCl, 25 NaHCO_3_, 3 KCl, 2 CaCl_2_, 2 MgSO_4_, 1 NaH_2_PO_4_, 0.5 Na-ascorbate, osmolarity adjusted to 310 mOsm with dextrose, pH 7.35) (Miska et al., 2018), the 300 μm slices were immediately transferred to a warm (34°C) chamber filled with a continuously carbogenated choline-based solution (in mM: 110 Choline-Cl, 25 NaHCO_3_, 11.6 Na-ascorbate, 7 MgCl_2_, 3.1 Na-pyruvate, 2.5 KCl, 1.25 NaH_2_PO_4_, and 0.5 CaCl_2_, adjusted to 310 mOsm with dextrose, pH 7.35) (Ting et al., 2014). After 5 min, slices were then transferred to warm (34°C) carbogenated standard ACSF and incubated another 30 min before being moved to room temperature. Slices were used for electrophysiology within 6 hours of slicing.

### CaMPARI2 *ex vivo* photoconversion

For *ex vivo* photoconversion experiments, 300 μm slices were prepared as above. After incubation, a slice was placed on the recording rig in active ACSF (in mM: 126 NaCl, 25 NaHCO_3_, 3.5 KCl, 1 CaCl_2_, 0.5 MgCl_2_, 1 NaH_2_PO_4_, 0.5 Na-ascorbate, osmolarity adjusted to 310 mOsm with dextrose, pH 7.35) (Maffei et al., 2004), then uniformly illuminated through a Blue Fluorescent Protein filter cube (TLV-U-MF2-BFP, Thorlabs) at ~0.20 mW light (measured at 390 mn, the peak excitation frequency of the filter cube) for 30 minutes. The uniformity of illumination allowed us to compare the red/green ratio between all neurons at the same slice depth. Active ACSF was replaced with regular ACSF before starting the recordings.

### Immunostaining

Animals used for immunostaining were deeply anesthetized with a ketamine/xylazine/acepromazine solution following photoconversion, then transcardially perfused with chilled 4% paraformaldehyde (PFA) in 0.01M PBS. Brains were removed, then post-fixed in PFA overnight before slicing into 50μm coronal slices the following day. Slices were washed three times before being stored in PBS with 0.05% NaN_3_ at 4°C.

Slices were incubated in a blocking and primary antibody solution (0.3% TritonX-100, 0.05% NaN_3_, 1% BSA, anti-cFos (9F6, Cell Signaling #2250, rabbit, 1:1000) in 0.01M PBS) for 24 hours. Anti-redCaMPARI2 (mouse, 1:1000, Janelia Research Campus) was then added and sliced were incubated overnight. The following day, slices were rinsed three times in PBS, then incubated for 2 hours in a secondary antibody solution (1% BSA, AlexaFluor goat anti-rabbit 647 (Invitrogen, 1:300), AlexaFluor goat anti-mouse 555 (Invitrogen, 1:300) in 0.01M PBS). Slices were then rinsed three times, then mounted and coverslipped using Fluoromount-G (SouthernBiotech). Slides were allowed to dry overnight before imaging with 488 nm, 543 nm, and, 647 nm lasers on a Leica SP5 confocal microscope.

### Electrophysiology

For *in vivo* and *ex vivo* experiments, images were captured in the red and green channels (mCherry and GFP filter cubes) before each experiment using a Hamamatsu C4742-95-12ERG camera. Neurons were visualized on an Olympus BX51QWI upright epiflouresence microscope with a 10× air (0.13 NA) and 40× water immersion objectives (0.8 NA) with infrared differential interference contrast (DIC) optics and an infrared CCD camera. V1m was identified by the shape and morphology of the white matter. In initial experiments using a pan-neuronal promoter, excitatory neurons were identified by their apical dendrite and teardrop shaped soma; neurons were filled with biocytin for post hoc reconstruction. All recordings were performed on slices continuously perfused with carbogenated 34 °C ACSF. Blockers and variations in ACSF composition are described below. Data were low-pass filtered at 6 kHz and acquired at 10 kHz with a Multiclamp 700B amplifier and a CV-7b headstage (Molecular Devices). Data were acquired using WaveSurfer (v0.953, Janelia Research Campus, Louden County, VA), and were analyzed using custom MATLAB scripts (Hengen et al., 2013; Lambo and Turrigiano, 2013; Miska et al., 2018). Potentials were not adjusted for the liquid junction potential. *In vivo* firing rate data in Figure 2D from Hengen et al., 2016 were reanalyzed by layer position of electrode, and plotted separately for all layers or for units from layer 4 only. Data were recording using WaveSurfer v0.953 (Janelia Research Campus)

#### Spontaneous firing measurements

To measure spontaneous firing, neurons were recorded in active ACSF (see above). Pipettes with a 2-5 mΩ resistance were filled with active ACSF, then used to form a loose seal (10-50 mΩ) with the targeted neuron (Maffei et al., 2004). Spontaneous activity was then recorded from pairs of nearby neurons (either simultaneously or sequentially) for 15 minutes. Neurons that did not fire any action potentials during the recorded period were discarded, as their ability to fire action potentials could not be verified. Thus, we are likely under sampling very low firing rate neurons with this procedure.

#### Intrinsic excitability measurements

To measure intrinsic excitability, we performed whole cell recordings with an internal recording solution containing (in mM) 100 K-gluconate, 10 KCl, 10 HEPES, 5.37 biocytin, 10 Na_2_-phosphocreatine, 4 Mg-ATP, and 0.3 Na-GTP, with sucrose added to bring osmolarity to 295 mOsm and KOH added to bring pH to 7.35 in pipettes with resistance between 4 and 8 mΩ (Lambo and Turrigiano, 2013; Miska et al., 2018). Synaptic currents were blocked by adding picrotoxin (PTX), 6,7-dinitroquinoxaline-2,3-dione (DNQX), and (2R)-amino-5-phosphonovaleric acid (APV) to standard ACSF to block γ-aminobutyric acid (GABA), α-amino-3-hydroxy-5-methyl-4-isoxazolepropionic acid (AMPA), and N-methyl-d-aspartate (NMDA) receptors, respectively. A small dc bias current was injected to maintain resting membrane potential around −70 mV, and 500 ms current injections of increasing amplitude ranging from −100 pA to 400 pA in intervals of 20 pA were delivered every 4 s (Desai et al., 1999). Neurons were excluded from analysis if they displayed Rs > 25 mΩ, Vm > −50 mV, or Rin < 80 mΩ. Action potential timings and waveforms were quantified with custom MATLAB scripts (Mathworks).

#### Synaptic input measurements

To measure excitatory and inhibitory synaptic charge, we performed whole cell recordings in active ACSF with an internal recording solution containing (in mM) 115 Cs-methanesulfonate, 10 HEPES, 10 BAPTA.4Cs, 5.37 biocytin, 2 QX-314 Cl, 1.5 MgCl2, 1 EGTA, 10 Na_2_-Phosphocreatine, 4 ATP-Mg, and 0.3 GTP-Na, with sucrose added to bring osmolarity to 295 mOsm and CsOH added to bring pH to 7.35; patch pipettes had resistances between 4 and 8 mΩ (Miska et al., 2018). In voltage clamp, the reversal potential for inhibition was determined by holding the cells at −50, and decreasing the holding potential in 5 mV increments until outward currents were not detectable. The procedure was then repeated for excitation, starting at −10 mV and increasing in 5 mV increments. Cells were then held at the experimentally determined reversal potential for inhibition (typically −65 mV to −50 mV) for 2 minutes, then switched to the experimentally determined reversal potential for excitation (typically −5mV to +10 mV) for 2 minutes; three independent measurements at each potential were made for each neuron, and averaged.

### Statistical analysis

Data analysis was performed using in house scripts in MATLAB, or using Graph Pad Prism (Hengen et al., 2013; Miska et al., 2018). The number of animals and neurons for each experiment, as well as statistical tests used, is given in the figure legends. Individual data points indicate measurements from single neurons. Error bars represent SEM. Data were tested for normality using a Shapiro Wilk normality test, and compared with a paired t-test (for paired, normally distributed data), Wilcoxon signed rank sum test (paired, non-normally distributed data), Mann Whitney U test (non-paired, non-normally distributed data), or Spearman’s rank correlation (non-normally distributed regression analysis).

## Acknowledgements

We thank B. Moeyaert and E. Schreiter for their gift of the CaMPARI2 viral construct. We thank Alejandro Torrado Pacheco for providing layer information for the *in vivo* firing rate data from V1. Supported by NS R35111562 (GGT) and NS F32101832 (NFT).

## Supplemental Material

**Figure S1: Input resistance is correlated with area under the F-I curve and rheobase. A**: Correlation between input resistance and area under F-I curve (for 10 AP current step) for all neurons. B: Correlation between input resistance r = Spearman’s rank correlation coefficient. All recordings and images are from pyramidal neurons in L4 V1.and rheobase current (for 10 AP current step) for all neurons.

**Table 1: Additional electrophysiological properties of high and low activity neurons.** Values labeled “rheobase” are calculated for the first spike at rheobase, values labeled “5 spikes” are calculated for the first spike for the smallest current step to produce at least 5 action potentials. Passive properties are measured in voltage clamp with a −5mV voltage step.

