## Supplementary figures and images for "Activity labeling *in vivo* using CaMPARI2 reveals electrophysiological differences between neurons with high and low firing rate set points"

### Supplemental Figure 1

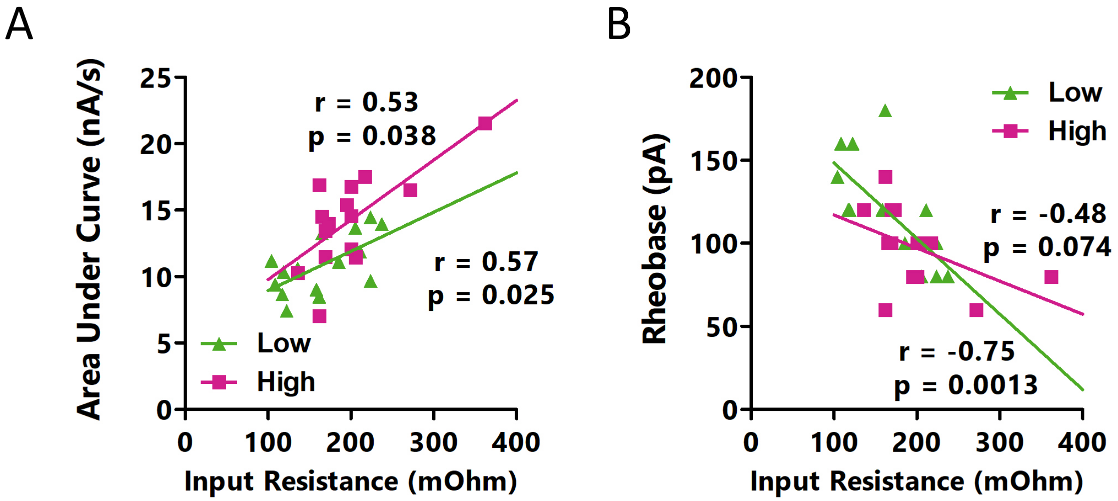
