## Supplemental Table 1 for "Activity labeling *in vivo* using CaMPARI2 reveals electrophysiological differences between neurons with high and low firing rate set points"

| | Parameter | | Median | Mean | Std. Dev | 95% CI | | p value |
|---|---|---|---|---|---|---|---|---|
| Rheobase | Spike amplitude (mV) | Low | 61.64 | 60.86 | 9.97 | 55.34 | 66.38 | 0.723 |
| | | High | 59.90 | 59.93 | 8.53 | 55.20 | 64.65 | |
| | AHP amplitude (mV) | Low | 27.98 | 29.22 | 7.19 | 25.24 | 33.21 | 0.941 |
| | | High | 28.30 | 29.05 | 6.75 | 25.31 | 32.79 | |
| | Mean rise slope (V/s) | Low | 152.90 | 150.20 | 23.35 | 137.30 | 163.10 | 0.719 |
| | | High | 149.60 | 147.40 | 29.14 | 131.30 | 163.60 | |
| | Mean decay slope (V/s) ^ | Low | 67.98 | 66.00 | 11.20 | 59.80 | 72.20 | 0.679 |
| | | High | 67.10 | 65.67 | 7.42 | 61.56 | 69.78 | |
| | Half width (ms) | Low | 0.30 | 0.35 | 0.07 | 0.31 | 0.39 | 0.865 |
| | | High | 0.30 | 0.35 | 0.12 | 0.29 | 0.42 | |
| | Maximum rise slope (V/s) | Low | 328.70 | 330.30 | 47.23 | 304.20 | 356.50 | 0.935 |
| | | High | 329.10 | 329.30 | 49.84 | 301.70 | 356.90 | |
| | Maximum decay slope (V/s) ^ | Low | 91.06 | 89.55 | 13.71 | 81.95 | 97.14 | 0.847 |
| | | High | 90.73 | 90.42 | 11.56 | 84.02 | 96.83 | |
| | Max rise/Max decay | Low | 3.61 | 3.72 | 0.38 | 3.51 | 3.93 | 0.775 |
| | | High | 3.55 | 3.67 | 0.50 | 3.39 | 3.95 | |
| | Latency (ms) ^ | Low | 14.46 | 16.01 | 7.68 | 11.76 | 20.26 | 0.169 |
| | | High | 10.47 | 12.59 | 6.08 | 9.22 | 15.95 | |
| 5 spikes | Spike amplitude (mV) | Low | 55.56 | 57.70 | 8.61 | 52.93 | 62.47 | 0.678 |
| | | High | 57.63 | 58.77 | 8.35 | 54.15 | 63.40 | |
| | AHP amplitude (mV) | Low | 29.78 | 30.34 | 6.56 | 26.71 | 33.97 | 0.680 |
| | | High | 29.36 | 29.35 | 6.13 | 25.95 | 32.74 | |
| | Mean rise slope (V/s) | Low | 138.90 | 143.20 | 23.89 | 130.00 | 156.50 | 0.895 |
| | | High | 151.80 | 142.20 | 26.00 | 127.80 | 156.60 | |
| | Mean decay slope (V/s) | Low | 63.94 | 60.20 | 12.03 | 53.53 | 66.86 | 0.804 |
| | | High | 59.67 | 59.48 | 8.10 | 55.00 | 63.96 | |
| | Half width (ms) | Low | 0.40 | 0.40 | 0.11 | 0.34 | 0.46 | 0.865 |
| | | High | 0.40 | 0.41 | 0.12 | 0.34 | 0.47 | |
| | Maximum rise slope (V/s) | Low | 317.20 | 314.30 | 52.78 | 285.00 | 343.50 | 0.835 |
| | | High | 322.50 | 317.40 | 46.01 | 291.90 | 342.80 | |
| | Maximum decay slope (V/s) ^ | Low | 89.42 | 84.57 | 15.88 | 75.78 | 93.37 | 0.978 |
| | | High | 88.76 | 84.35 | 12.05 | 77.68 | 91.03 | |
| | Max rise/Max decay | Low | 3.83 | 3.77 | 0.58 | 3.45 | 4.10 | 0.891 |
| | | High | 3.80 | 3.80 | 0.52 | 3.51 | 4.09 | |
| | Latency (ms) | Low | 6.95 | 6.97 | 2.25 | 5.72 | 8.22 | 0.830 |
| | | High | 7.25 | 7.15 | 2.29 | 5.88 | 8.41 | |
| | Input resistance (mΩ) ^ | Low | 161.80 | 165.30 | 46.58 | 139.50 | 191.10 | 0.079 |
| | | High | 195.00 | 199.20 | 55.20 | 168.60 | 229.70 | |
| | Capacitance (pF) ^ | Low | 75.59 | 78.25 | 11.56 | 71.85 | 84.65 | 0.083 |
| | | High | 70.41 | 69.66 | 13.40 | 62.24 | 77.08 | |
| | Resting membrane potential (mV) | Low | -66.70 | -66.86 | 4.14 | -69.15 | -64.57 | 0.572 |
| | | High | -67.45 | -67.40 | 3.58 | -69.39 | -65.42 | |
| | Action potentail threshold (mV) ^ | Low | -30.85 | -31.47 | 2.82 | -33.03 | -29.91 | 0.208 |
| | | High | -32.83 | -33.14 | 4.93 | -35.87 | -30.41 | |
| | Sag percentage ^ | Low | 94.04 | 90.31 | 9.61 | 84.99 | 95.63 | 0.252 |
| | | High | 94.67 | 93.73 | 4.04 | 91.49 | 95.97 | |

p values calculated via paired t-test except ^, which were calculated via Wilcoxon Signed Rank Sum test
